# Rapid FFR: A rapid method for obtaining Frequency Following Responses

**DOI:** 10.1101/2025.05.20.655073

**Authors:** Jonas Huber, Tim Schoof, Hannah Stewart, Alain de Cheveigné, Stuart Rosen

## Abstract

**Objectives:** Frequency-Following Responses (FFRs) are typically recorded using stimuli of 40–250 ms in duration separated by silent intervals. Because robust FFRs require averaging across approximately 3000 stimulus repetitions, conventional acquisition is time-intensive. We introduce the Rapid FFR, a technique that minimises recording time by presenting the stimulus continuously (i.e., without inter-stimulus intervals) and by averaging across individual response cycles. This study aimed to: (a) compare Rapid and Conventional FFR performance under time-matched conditions; (b) assess test-retest reliability of both methods, and (c) examine potential neural adaptation during extended Rapid FFR recordings.

**Design:** All participants (37 in total) were young adults with clinically normal hearing. In Experiment 1, FFRs were elicited from 16 listeners using a 128 Hz sawtooth wave presented in two ways: (1) continuously over approximately 1 minute in each polarity for the Rapid FFR and (2) as discrete tone bursts (1500 trials per polarity) for the standard FFR. Signal-to-Noise ratios (SNRs), response amplitudes, and test–retest reliability were compared across techniques. In Experiment 2, the Rapid FFR was recorded continuously from 21 listeners for nearly nine minutes to assess potential adaptation over time.

**Results:** The Rapid FFR produced significantly higher SNRs than the standard FFR, reflecting improved recording efficiency. Both techniques captured inter-individual differences reliably, with comparable frequency-domain response patterns across participants. In particular, measures derived from the Rapid FFR showed strong agreement with those from the standard FFR for lower harmonics (F0–H3). Higher harmonics (H4–H7) exhibited greater variability but remained consistent between techniques. In the prolonged recording condition, FFR amplitudes remained stable over time, with no significant decline across the nine minute recording. This indicates that continuous stimulation did not result in measurable neural adaptation or response fatigue.

**Conclusions:** The Rapid FFR offers a time-efficient alternative to standard protocols, preserving signal fidelity and reliability while enabling shorter acquisition times. These findings suggest that the Rapid FFR is well-suited for use in populations where long recordings are challenging (e.g., in infants and the clinic) and can facilitate more extensive experimental designs within a single session. The method holds promise for advancing both research and clinical applications of the FFR. Future studies should explore its use across a broader range of sounds (in particular, dynamically varying ones) and listener groups.

## 1 Introduction

The Frequency Following Response (FFR) is a scalp-recorded electrophysiological measure that reflects sustained synchronous neural firing in the brainstem in response to the auditory presentation of periodic stimuli (Marsh & Worden, 1968; Moushegian et al., 1973). FFRs are typically recorded in response to a stimulus presented in both positive and negative polarities, and the responses to both polarities are then either added or subtracted. The summed-polarity FFR, often referred to as the Envelope Following Response (EFR), is believed to eliminate linear stimulus artifacts and the cochlear microphonic (Aiken & Picton, 2008), thereby emphasising the envelope of the response (Gorga et al., 1985). Conversely, the subtracted-polarity FFR, also known as the Temporal Fine Structure (TFS), emphasises higher-frequency components (Aiken & Picton, 2008).

Typically, FFRs are recorded using a repeated stimulus of 40–250 ms duration, sometimes extending up to two seconds with interstimulus intervals of around 50 ms (Patel et al., 2023; Fuh-Cherng Jeng & Li, 2011; Greenberg et al., 1987; Dajani et al., 2005). Robust FFR recordings require significant time, as stimuli are typically presented approximately 1500 times each in the two polarities, resulting in sessions lasting up to 15 minutes (Patel et al., 2023; Skoe, Burakiewicz, Figueiredo, & Hardin, 2017). Here, we introduce an alternative technique, the **Rapid FFR** to accelerate data acquisition by presenting stimuli continuously (without interstimulus intervals) and averaging across a single cycle of the response.

The use of prolonged auditory stimulation in EEG recordings is not entirely novel. For example, the auditory steady-state response, a cortical electrophysiological measure elicited by amplitude-or frequency-modulated tones, is typically recorded with continuous stimulus presentation (e.g., Cone-Wesson et al., 2002; Dimitrijevic et al., 2002). Continuous stimulus presentation has also been sporadically used for FFR recording (Batra et al., 1986; Aiken & Picton, 2008). For instance, Aiken and Picton (2008) recorded FFRs in response to 1.5-second-long natural vowels continuously presented for 75 seconds (50 repetitions).

The Rapid FFR further innovates by averaging across a single cycle of the response, which is typically very short in steady-state periodic sounds, being 1/fundamental frequency (F0) long (e.g., 10 ms for an F0 = 100 Hz). In Conventional FFR experiments using steady-state sounds, averaging is done over several cycles (see for sine waves in Gorina-Careta, Kurkela, Hämäläinen, Astikainen, & Escera, 2021 or harmonic complexes in Krishnan, 2002). Averaging over a single cycle in the Rapid FFR increases the effective number of trials, thereby attenuating noise in the resulting response waveform more efficiently.

In combining continuous stimulation with single-cycle averaging, we build on previous research that already did so in sinusoids (only very recently discovered by us - D. Parker & Matsebula, 1998; D. J. Parker & Matsebula, 1992), now extending this approach to complex steady-state sounds.

The primary aim of this study is to determine if the Rapid FFR allows for more time-efficient data collection than the Conventional FFR protocol. Secondly, we aim to demonstrate that the Rapid and Conventional FFR techniques yield similar results for both the summed-and subtracted polarity FFR. We evaluate this by investigating whether the Rapid FFR is equally sensitive to inter-participant differences as the Conventional FFR. Finally, we investigate the extent of neural adaptation during extended recording periods of the Rapid FFR, an important factor in determining its benefits relative to the conventional method (G. Bidelman & Powers, 2018; Gorina-Careta, Zarnowiec, Costa-Faidella, & Escera, 2016).

Two experiments address these questions. Experiment 1 examines the relationship between the Rapid FFR and the Conventional FFR, while Experiment 2 investigates the potential effects of neural adaptation within the Rapid FFR over a longer time window (≈ 9 minutes).

## 2 Materials and Methods

### 2.1 Participants

Two independent samples of 16 and 21 young listeners, aged 20–29 years (9 female) and 18–30 years (13 female), participated in Experiments 1 and 2, respectively. All participants had normal hearing, defined as pure-tone thresholds of 20 dB HL or better at octave frequencies between 250 and 8000 Hz, and normal click ABRs (see Appendix for more details). None of the participants reported a history of neurological disorders. They were compensated for their participation and provided informed consent, which was approved by the UCL Research Ethics Committee.

### 2.2 Stimuli

FFRs were recorded in response to a harmonic complex (sawtooth wave) with a fundamental frequency (F0) of 128 Hz, generated by the additive synthesis of 30 harmonics. The F0 was chosen as a divisor of the sampling rate used for FFR collection (16384 Hz), ensuring that the timing of each F0 cycle in the continuous EEG signal could be determined with a precision of one sample point. To avoid transients in the audio signal, the stimulus was always tapered on and off over one cycle, i.e. 7.8125 ms.

In Experiment 1, the stimulus was presented in two ways. In the conventional technique, each stimulus lasted 54.7 ms (equivalent to seven F0 cycles), with a 45 ms interstimulus interval between repetitions. A total of 3000 repetitions were presented in two blocks: one block containing 1500 positive-polarity trials and the other block containing 1500 negative-polarity trials.

The Rapid FFR technique involved presenting 7503 cycles (approximately 58 seconds) of the sawtooth wave for each polarity, with the two polarities separated into distinct blocks. In both the rapid and conventional technique, blocks were separated by a brief interval of silence.

In Experiment 2, the sawtooth wave had a duration of 67500 F0 cycles (approximately 9 minutes) in each polarity. As in Experiment 1, the two polarities were presented in individual blocks, each separated by a brief interval of silence.

### 2.3 Procedure

Participants were seated in a reclining chair within an electrically shielded soundproof booth and were allowed to fall asleep. FFRs were collected using a BioSemi ActiveTwo system (Amsterdam, The Netherlands). FFRs were recorded from the top of the scalp (Cz) and referenced against the seventh cervical vertebra, situated at the back of their neck. Two additional electrodes, Common Mode Sense (CMS) and Driven Right Leg (DRL), were used in place of a traditional ground electrode in the BioSemi ActiveTwo system. Electrode offsets were consistently less than 40 µV, and responses were recorded with a sampling rate of 16384 Hz.

The stimuli were generated using MATLAB (MathWorks, Natick, MA) with stimuli on one channel and triggers on a separate channel. Both channels were delivered via a computer equipped with an external sound card (RME FireFace UC) and connected to a custom-made trigger box. This trigger box separated the two channels, sending the stimulus to electrically shielded insert earphones (ER-3, Intelligent Hearing Systems, Miami, FL) and the trigger to the EEG system. Stimuli were presented binaurally at a level of 80 dB SPL.

In Experiment 1, responses were recorded using the two different methods, as detailed in Section 2.2. Each recording technique was used twice to assess test-retest reliability, and the presentation order of conditions was balanced across participants. In Experiment 2, only the rapid technique was used, and FFRs were measured just once.

## 3 Analysis

Data preprocessing was performed using custom scripts in MATLAB and Python. In MATLAB, the EEGLAB toolbox (A & S, 2004) was employed, while in Python, the `numpy` (Harris et al., 2020), `pandas` (The Pandas Development Team, 2020), `maths` (Van Rossum, 2020), and `concurrent` (Python Core Team, 2019) libraries were used.

### 3.1 Preprocessing Rapid FFR

For each combination of participant and polarity, we selected 7503 cycles in experiment 1 and 67500 cycles in experiment 2 of EEG data, starting 10 ms after stimulus onset to account for the delay in the signal reaching the brainstem (Picton et al., 2003). The data vectors were then resampled to correct for the sampling asynchrony between the presenting computer and the BioSemi system, which resulted from slightly different internal clock frequencies. The resampled data vectors were divided into individual trials, each consisting of 128 data points (7.8125 ms) in length. Trials containing values deviating more than 35 µV from 0 were rejected as part of the artifact rejection process. On average, 9.1 trials per polarity were excluded in experiment 1, and 396.7 in experiment 2. The remaining trials were averaged to form an Event-Related potential (ERP). The ERPs for positive and negative polarities were then added or subtracted to obtain the Envelope-Frequency Response (EFR) and Temporal Fine Structure (TFS), respectively (Aiken & Picton, 2008).

We did not zero-pad the waveform, as the frequency bins were already perfectly aligned with the frequencies of interest. Fourier analyses on the EFR and TFS were used to calculate the root mean square (RMS) levels of the harmonics. The values from the first three harmonics (128 Hz, 256 Hz, and 384 Hz) were derived from the EFR and corresponded to their respective frequency bins. Similarly, the RMS levels for the last four harmonics (512 Hz, 640 Hz, 768 Hz, and 896 Hz) were TFS-derived and matched their respective frequency bins.

Of primary interest in this study are the signal-to-noise ratios obtained from the two different methods. In typical FFR testing, the level of the noise is estimated either from the silent periods between stimuli, or from energy in the spectrum between the harmonics. Neither of these is available with the Rapid FFR method specified here. Therefore, to estimate noise levels, we combined bootstrapping with permutation testing on the resampled and artifact-excluded data vector (described above) from each participant and each polarity. Namely, a random starting point was selected within the vector, ranging from one to the total number of data points minus the length of one cycle. From this starting point, we extracted 128 data points (one cycle lasting 7.8125 ms) and labeled it as one noise trial. This procedure was repeated as many times as there were trials in the original dataset. All noise trials were averaged to form noise-ERPs, and the subsequent calculations for noise-EFR and noise-TFS followed the same procedures described for the signal.

The rationale behind this process is that by randomly selecting start points in the data vector, phase-locked signals are effectively canceled out due to phase cancellation, resulting in a noise spectrum devoid of phase-locked activity.

This procedure was repeated 10000 times, yielding 10000 noise-EFR and noise-TFS spectra. The “true” noise values were then calculated as the medians of these 10000 noise-EFR and noise-TFS spectra in each harmonic frequency bin. For example, the true noise value for one participant at F0 was the median of the 10000 noise-EFR RMS values at the 128-Hz-bin. SNRs were later calculated by contrasting the signal values with the true noise values in each harmonic frequency bin using the formula:

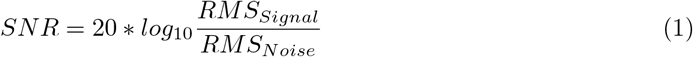

### 3.2 Preprocessing Conventional FFR

The analysis of Conventional FFR data involved selecting 3000 data segments (1500 per polarity), each 54.7 ms long (referred to as “trials”), starting at a specific time after stimulus onset. This time lag was determined by identifying the point between 5 and 15 ms after stimulus onset where the cross-correlation of the EFR with a 128 Hz sine wave was maximal.

The data segments were resampled as described in Section 3.1 and baseline-corrected using a 7 ms interval before stimulus onset. Trials with values deviating more than 35 µV from 0 were rejected as part of the artifact rejection process, resulting in an average of 8.0 rejected trials per polarity. The remaining trials were averaged to create two ERPs, one for each polarity.

**Figure 1.**
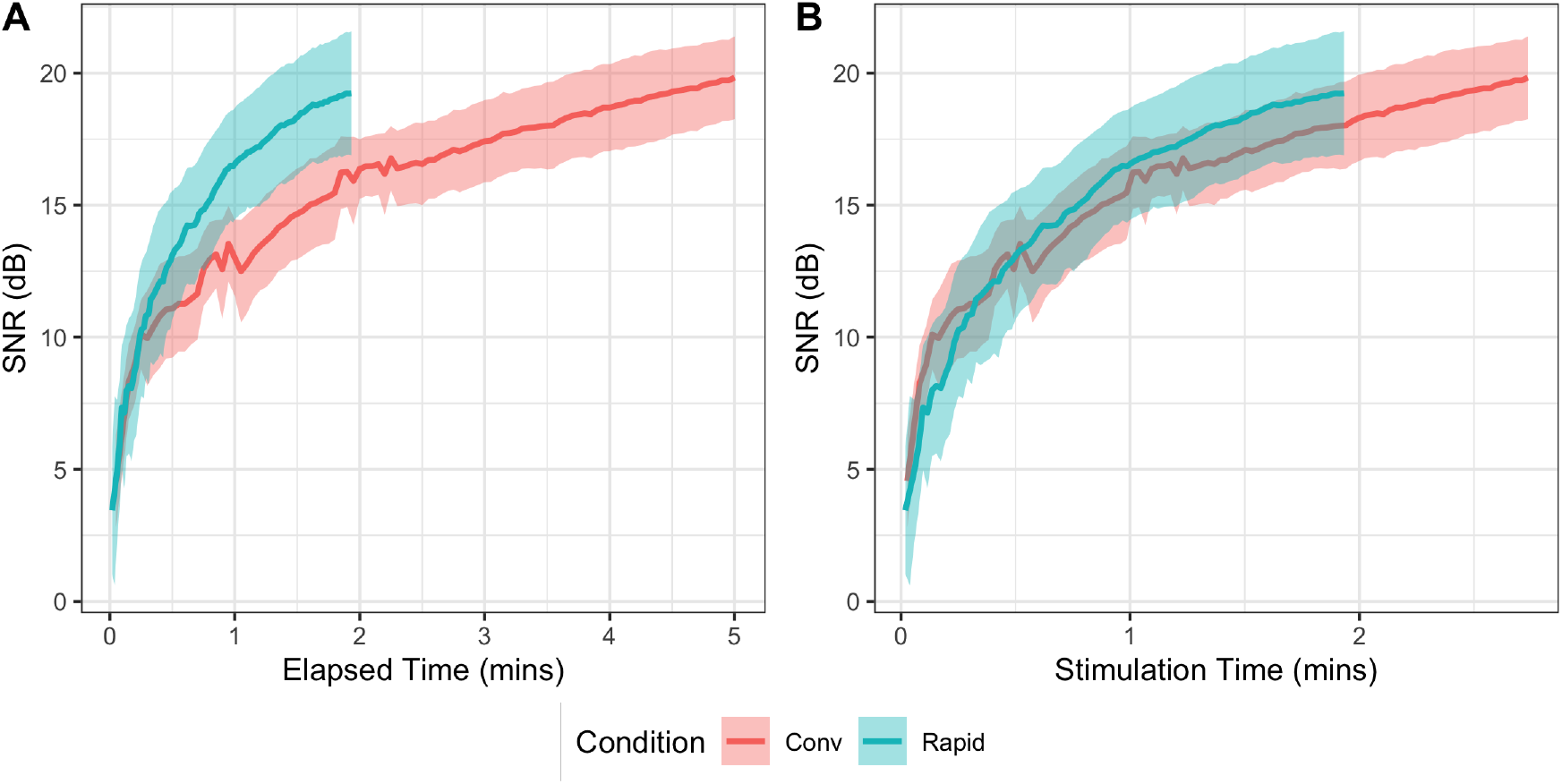
(A) The growth in SNR for the fundamental component (F0) of the recorded wave in the Rapid and Conventional FFR as a function of elapsed time, the actual time taken for the recording, including any silent periods. Lines represent the mean across participants, and shaded areas show the 95 % confidence intervals. Note that here and in all figures and analyses, the first 3 components are extracted from added polarity waveforms (the EFR), and the next 4 components from the subtracted polarities (the TFS). (B) The growth in SNR for the fundamental component (F0) of the recorded wave in the Rapid and Conventional FFR as a function of *stimulation time*, the total time for which there is a sound playing, thus excluding silences. Lines represent the mean across participants, and shaded areas show the 95 % Confidence intervals.

EFR and TFS ERPs were computed, again not zero-padding the waveform, as the frequency bins were already perfectly aligned with the frequencies of interest. Fourier analysis was then performed on the EFR and TFS data to obtain the RMS levels of the harmonics. As for the Rapid FFR, signal values for the first three harmonics were extracted from the EFR, and for harmonics 4-7, from the TFS. SNRs were estimated using the same method as described in Section 3.1, after concatenating together the data segments and with adjustments for trial length.

## 4 Results for Experiment 1 - Rapid vs Conventional FFR

The rest of the analysis was done in R, using the packages `tidyverse` (Wickham et al., 2019), `metap` (Dewey, 2022), `afex` (Singmann, Bolker, Westfall, & Aust, 2016), `lme4` (Bates, Mächler, Bolker, & Walker, 2015), `broom` (Robinson, Hayes, & Couch, 2022), and `irr` (Gamer, Lemon, Fellows, & Singh, 2019). For all datasets analysed in this study, we applied outlier rejection by excluding SNR values or signal amplitudes that deviated by more than 2.5 standard deviations from the mean. As a result, approximately 1% of the data was excluded across all analyses in this study.

### 4.1 Comparing the Rapid FFR with elapsed-time-matched Conventional FFR

First, we investigated whether the Rapid and Conventional FFR differed when matched for elapsed testing time — that is, the total time required to acquire the response, including breaks in the Conventional technique. This comparison addressed the practical question of whether the Rapid FFR is more efficient than the conventional protocol. Here we used the full Rapid FFR dataset and compared it to one minute of elapsed time in the Conventional FFR dataset in each polarity. As the elapsed time includes silent intervals, only 39% of the Conventional FFR dataset is used in this comparison (a total of 1170 data segments, or 585 trials per polarity). As the outcome measure, we used SNR values, as these account for both the reduction in noise within the recording and the strength of the signal.

To compare the two techniques, we employed a linear mixed effects model with the outcome variable of SNR at the maximum elapsed time. The necessity for any transformations (e.g., logarithmic or quadratic trends across harmonics) were identified from visual inspection of the data, both here and in subsequent analyses. The current model included two fixed effects and their interaction: *Harmonic* (F0-H7), treated as a continuous variable with both slope and quadratic term, and *Condition* (Rapid vs. Conventional FFR). Additionally, we included a random intercept for participant. The model revealed that the Rapid FFR exhibits significantly higher SNR levels than the Conventional FFR consistently across all harmonics (p *<* .001). No significant interaction between *Harmonic* and *Condition* was observed, either in the slope (p = .091) or the quadratic term (p = .88). An illustration of the data is provided in Figure 2, and detailed results for this and all subsequent statistical models are reported in the Appendix.

**Figure 2.**
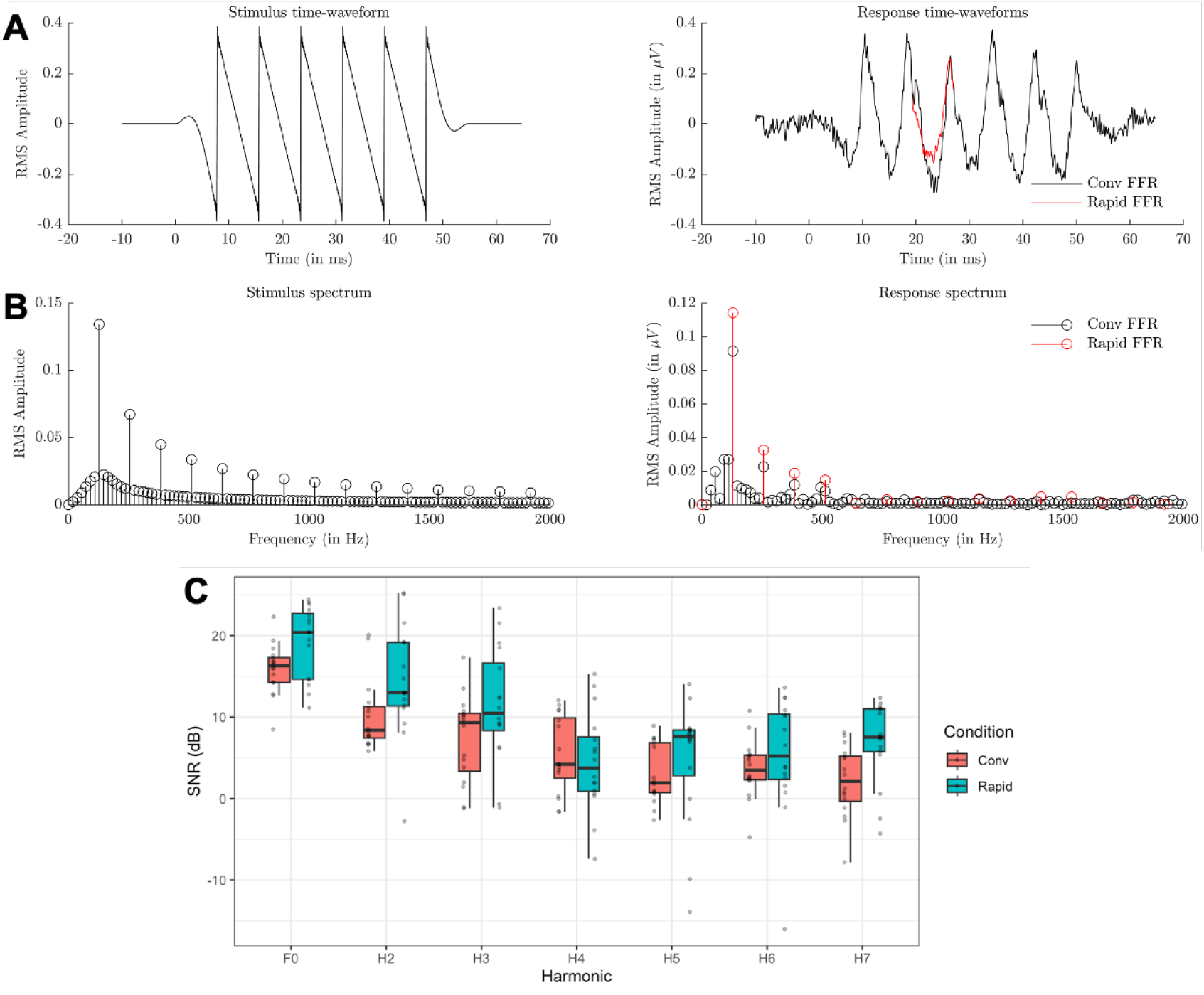
(A) Time-domain representation of stimulus (left) and EFR response (right), averaged across participants. The conventional FFR is shown in black, and the red line represents the 1-cycle average in the Rapid FFR condition. (B) Frequency-domain view of the same data. (C) Boxplot comparing Rapid vs Conventional FFR SNRs, with the conventional method cut to 1 minute of elapsed time for each polarity. Grey dots indicate individual datapoints.

To assess whether this improvement in SNR is solely attributable to the additional data, we performed an additional calculation. By artificially reducing the Rapid FFR dataset by 47% to match the data volume of the Conventional FFR, we observed that the two techniques produced statistically comparable SNRs. A linear mixed effects model with an identical structure to the previous analysis showed that the main effect of *Condition* was no longer significant (*p* = .116).

### 4.2 The relationship between the Rapid and the Conventional FFR measures

To determine whether the Rapid and Conventional FFR measure the same response, we compared the signal amplitudes between the two techniques, as signal amplitudes provide a more direct measure of the neural response strength, and because they are more widely used in the FFR literature than SNRs. For this analysis, we used the full dataset from both the Rapid and Conventional FFR, comprising 7603 trials for the Rapid FFR and 1500 trials for the Conventional FFR in each polarity, as this would ensure more accurate estimates of the response being measured.

We employed the intraclass correlation coefficient (ICC) using the R package irr. Briefly, the ICC is an extension of the common correlation coefficient that assesses the consistency or agreement between multiple measurements of the same unit. Unlike a standard correlation, the ICC can account for both systematic differences and random error, making it particularly useful in medical and research settings to evaluate reliability. We computed seven ICCs, one for each harmonic. The ICCs were based on a “oneway” model structure, which treats participants as random effects and the ‘raters’ (here methods, Rapid vs. Conventional FFR) as a fixed effect. The results are presented in Table 1, and the data are visualised in Figure 3.

**Table 1.** Intraclass correlation coefficient (ICCs) between the signal levels of the Rapid and Conventional FFR. All significant p-values remained significant after correction for multiple comparisons using the Bonferroni-Holm method, except for H6, which was corrected to a p-value of p = .093. Significant p-values here (and in following tables) are marked by a ‘*’.

| Harmonic | ICC-value | dF | 95% CI | uncorrected p-value |
| --- | --- | --- | --- | --- |
| F0 (EFR) | .664 | [15, 16] | [.279, .867] | .001 * |
| H2 (EFR) | .594 | [15, 16] | [.170, .835] | .005 * |
| H3 (EFR) | .650 | [15, 16] | [.257, .861] | .002 * |
| H4 (TFS) | .202 | [14, 15] | [-.315, .632] | .221 |
| H5 (TFS) | -.044 | [15, 16] | [-.505, .444] | .565 |
| H6 (TFS) | .493 | [14, 15] | [.009, .794] | .023 * |
| H7 (TFS) | -.217 | [15, 16] | [-.625, .292] | .800 |

**Figure 3.**
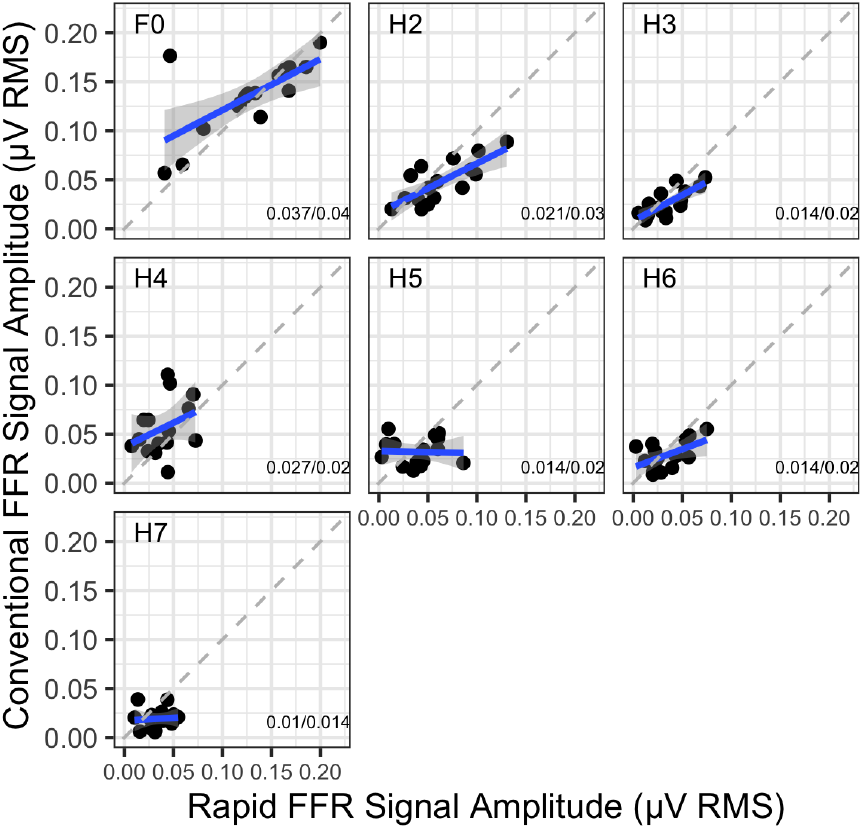
The relationship between the signal RMS levels measured in the Conventional and Rapid FFRs within participants. Here, we include the full datasets, i.e. 1 minute of elapsed time in the Rapid FFR in each polarity and 2.5 minutes of elapsed time in the Conventional FFR. On the bottom left corner, the SDs are shown in format *Conventional FFR SD/Rapid FFR SD*.

Interestingly, significant ICCs were observed for all 3 EFR harmonics, with ICCs exceeding .5. Among the higher TFS harmonics, only H6 demonstrated a similarly large value (ICC = .493, p = .023). Also, note in Figure 3 that the best-fit lines are always shallower than 1, reflecting at least partially a uniformly greater variability across participants for the Rapid than the Conventional technique.

### 4.3 Test-retest reliability in the Rapid and the Conventional FFR

Since the stimulus was presented twice in each condition, we also aimed to examine the test-retest reliability of the signal amplitudes in the Rapid FFR and compare them to those obtained with the Conventional FFR. Signal amplitudes were again chosen for this analysis. To ensure a realistic comparison of efficiency and reliability, we matched the Rapid and Conventional FFR datasets by elapsed time, and again calculated the ICC, but this time employed a “twoway” model structure, which treats both participants and raters (i.e., the first run vs. the second run) as random effects. The results are shown in Table 2, and the data are visualised in Figure 4.

**Table 2.**
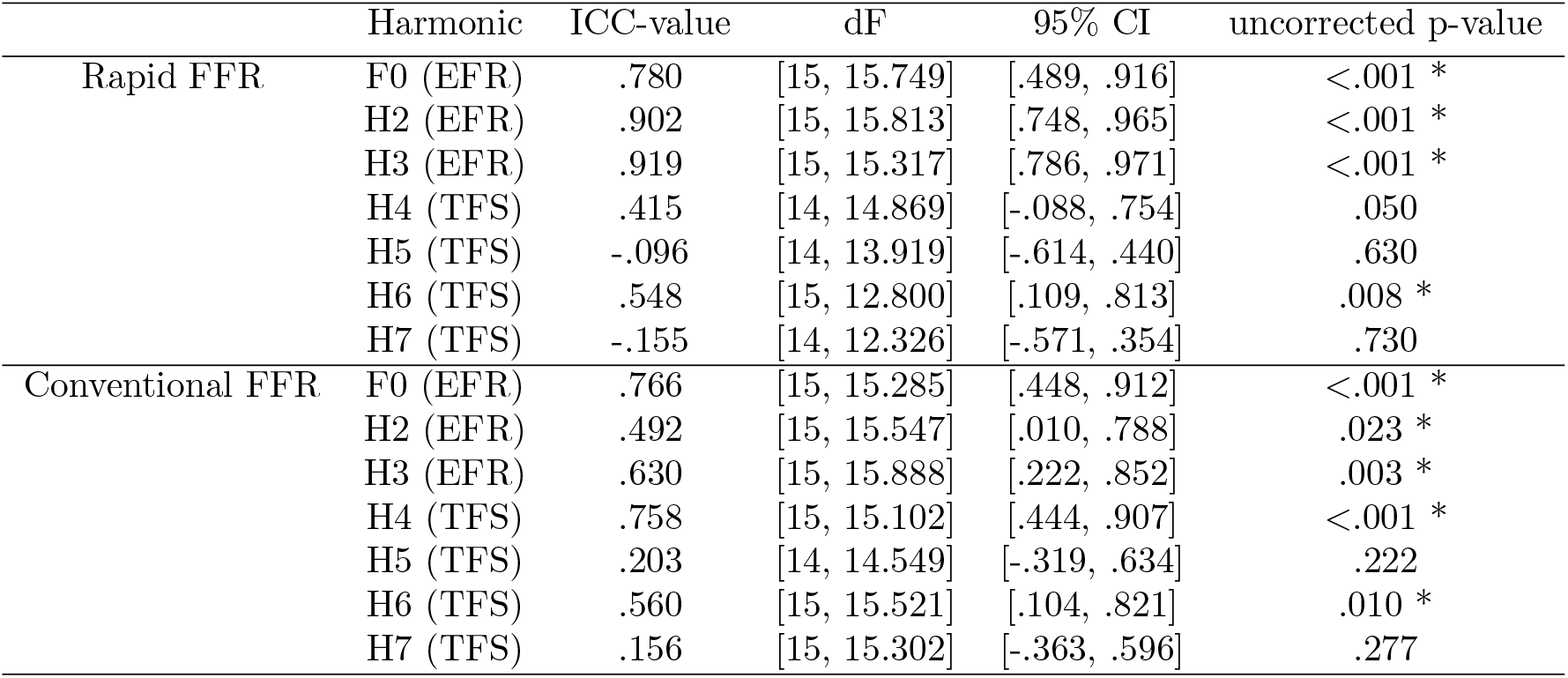
Results for the ICCs which correlate the signal levels between the two runs in each the Rapid FFR with Conventional FFR. p-values depicted here were not adjusted for multiple comparisons.

|  | Harmonic | ICC-value | dF | 95% CI | uncorrected p-value |
| --- | --- | --- | --- | --- | --- |
| Rapid FFR | F0 (EFR) | .780 | [15, 15.749] | [.489, .916] | <.001 * |
|  | H2 (EFR) | .902 | [15, 15.813] | [.748, .965] | <.001 * |
|  | H3 (EFR) | .919 | [15, 15.317] | [.786, .971] | <.001 * |
|  | H4 (TFS) | .415 | [14, 14.869] | [-.088, .754] | .050 |
|  | H5 (TFS) | -.096 | [14, 13.919] | [-.614, .440] | .630 |
|  | H6 (TFS) | .548 | [15, 12.800] | [.109, .813] | .008 * |
|  | H7 (TFS) | -.155 | [14, 12.326] | [-.571, .354] | .730 |
| Conventional FFR | F0 (EFR) | .766 | [15, 15.285] | [.448, .912] | <.001 * |
|  | H2 (EFR) | .492 | [15, 15.547] | [.010, .788] | .023 * |
|  | H3 (EFR) | .630 | [15, 15.888] | [.222, .852] | .003 * |
|  | H4 (TFS) | .758 | [15, 15.102] | [.444, .907] | <.001 * |
|  | H5 (TFS) | .203 | [14, 14.549] | [-.319, .634] | .222 |
|  | H6 (TFS) | .560 | [15, 15.521] | [.104, .821] | .010 * |
|  | H7 (TFS) | .156 | [15, 15.302] | [-.363, .596] | .277 |

**Figure 4.**
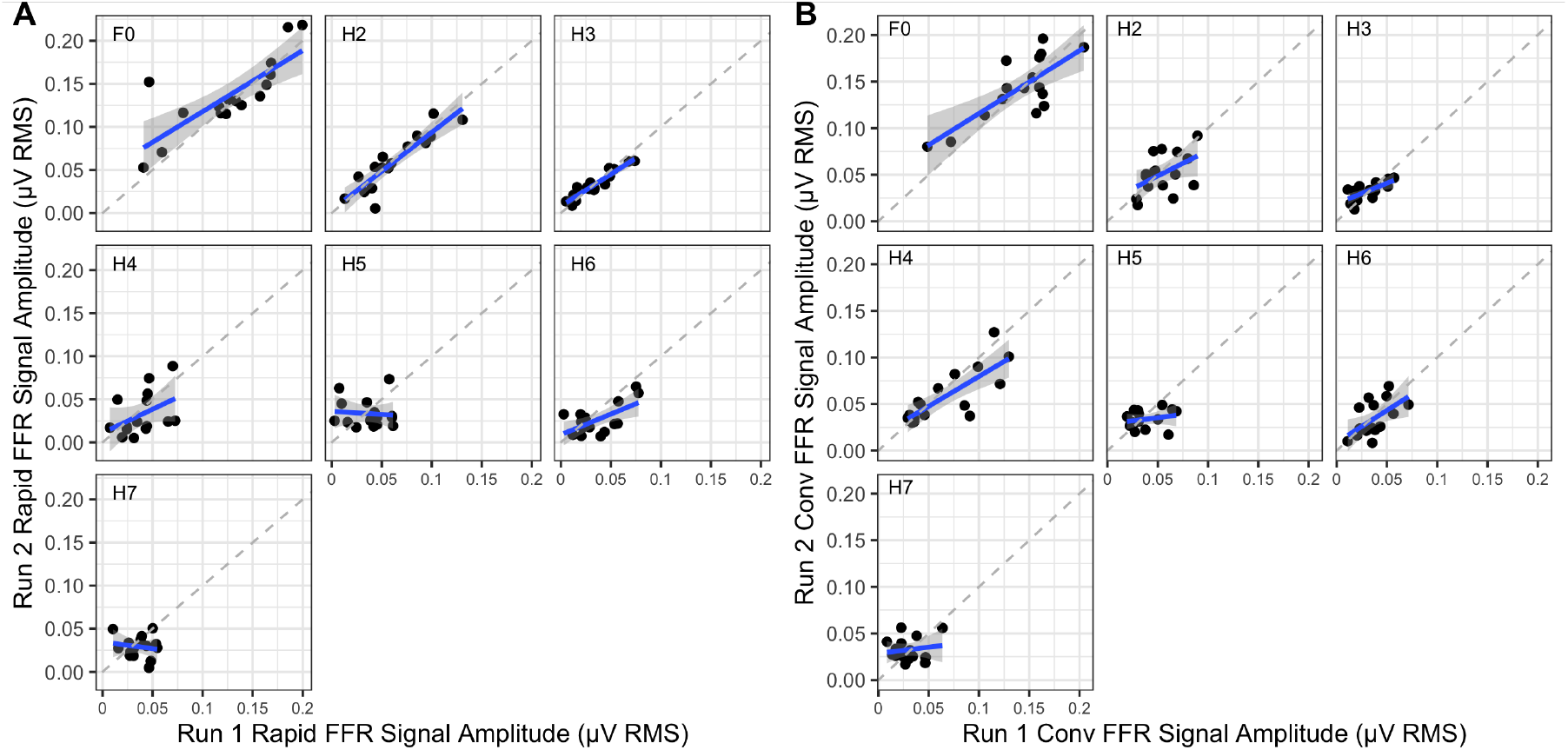
(A)The relationship between the signal RMS levels measured in the two runs of the Rapid FFR, again with elapsed time reduced to 1 minute per polarity. (B) The relationship between the signal RMS levels measured in the two runs of the Conventional FFR, with elapsed time reduced to 1 minute per polarity.

In the rapid technique, the analysis revealed significant ICCs for the lower three EFR harmonics, with remarkably high ICC values (ICC *>* .75). However, for the higher harmonics, the ICC values dropped significantly, and only H6 reached significance (ICC = .548, p = .008). Notably, all significant p-values shown in Table 2 in relation to the Rapid FFR remained significant after correction for multiple comparisons using the Bonferroni-Holm method.

In the conventional technique, the results were slightly more balanced across harmonics, with all harmonics except H5 and H7 showing significant ICCs, with values ranging between .20 and .77. However, after correcting for multiple comparisons using the Bonferroni-Holm method, only F0 (p = .001), H3 (p = .014), H4 (p *<* .001), and H6 (p = .040) retained significant p-values.

Lastly, we investigated whether the Rapid FFR exhibited statistically significantly different ICC values compared to the Conventional FFR. To assess this, we combined a bootstrapping method with permutation testing. Specifically, for the Rapid FFR at a single harmonic, we randomly matched the signal amplitudes from the first run with those of the second run, effectively scrambling participants’ scores. We then computed the ICC values of the resulting bootstrapped data and repeated this process for the Conventional FFR at the same harmonic. The ICC values of the Rapid and Conventional FFR were subtracted from each other, and this procedure was repeated 1000 times to generate noise ICC values for each frequency bin of interest. These values were used to establish the distribution of expected values under the null hypothesis, which is that there are no differences between the ICC values from the two methods. To estimate a p-value for the outcome statistic, we used the formula

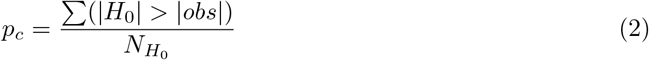

, in which *p*_*c*_ represents the p-value, *H*_0_ stands for the permuted noise values, *obs* denotes the observed difference between the actual ICC values for the Rapid and Conventional FFR, and *N*_*H*0_ indicates the number of noise values. Importantly, as this was a two-tailed test, we compared the absolute value of the observed ICC difference (|*obs*|) to the absolute values of the permuted noise values (|*H*_0_|). The p-value is therefore calculated as the proportion of absolute values in the null hypothesis distribution that are greater than the absolute observed ICC difference.

None of these p-values indicated statistical significance, with results of p = .955, .158, .312, .306, .386, .967, and .379 for F0 through H7, respectively. In short, there is no evidence that the two methods differ in their ICC values, demonstrating that the rapid technique — despite its faster acquisition — performs no worse in terms of test-retest reliability compared to the conventional FFR.

## 5 Results for Experiment 2 - Extended Duration Rapid FFR

In Experiment 2, we tested a different set of participants using the Rapid FFR technique only, presenting 67500 F0 cycles in each polarity (∼ 9 minutes). Our primary aim was to investigate the extent of neural adaptation during extended recording periods. We did this by segmenting our waves into 6 sequential *chunks*, each containing 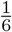 th of the data (90 s), which were then analysed independently. This choice was guided by the fact that a minimum duration of 90 s was required for all harmonics of interest to reach SNRs indicating a reliable signal, no longer dominated by noise.

We constructed four linear mixed effects statistical models, each using a different data set. These consisted of the SNRs and signal amplitudes (separately) for the F0 – H3 harmonics in the EFR and the H4-H7 harmonics in the TFS. All models followed the same overall structure. Each model included a fixed term for the continuous variable *Harmonic*, modelled as a quadratic function, and another fixed term for the continuous variable *Chunk*, also modeled as a quadratic function, along with a random intercepts for participant. After model simplification, only the model using the EFR SNR values retained significant fixed terms (p = .011 for the slope and p = .037 for the quadratic term) relating to *Chunk*. In contrast, the other three models — those for the TFS SNR values and the TFS and EFR signal amplitudes — retained no significant interactions or main effects involving *Chunk*. The results for the signal amplitudes are shown in Figure 5, while the figure displaying SNRs is provided in the Appendix.

**Figure 5.**
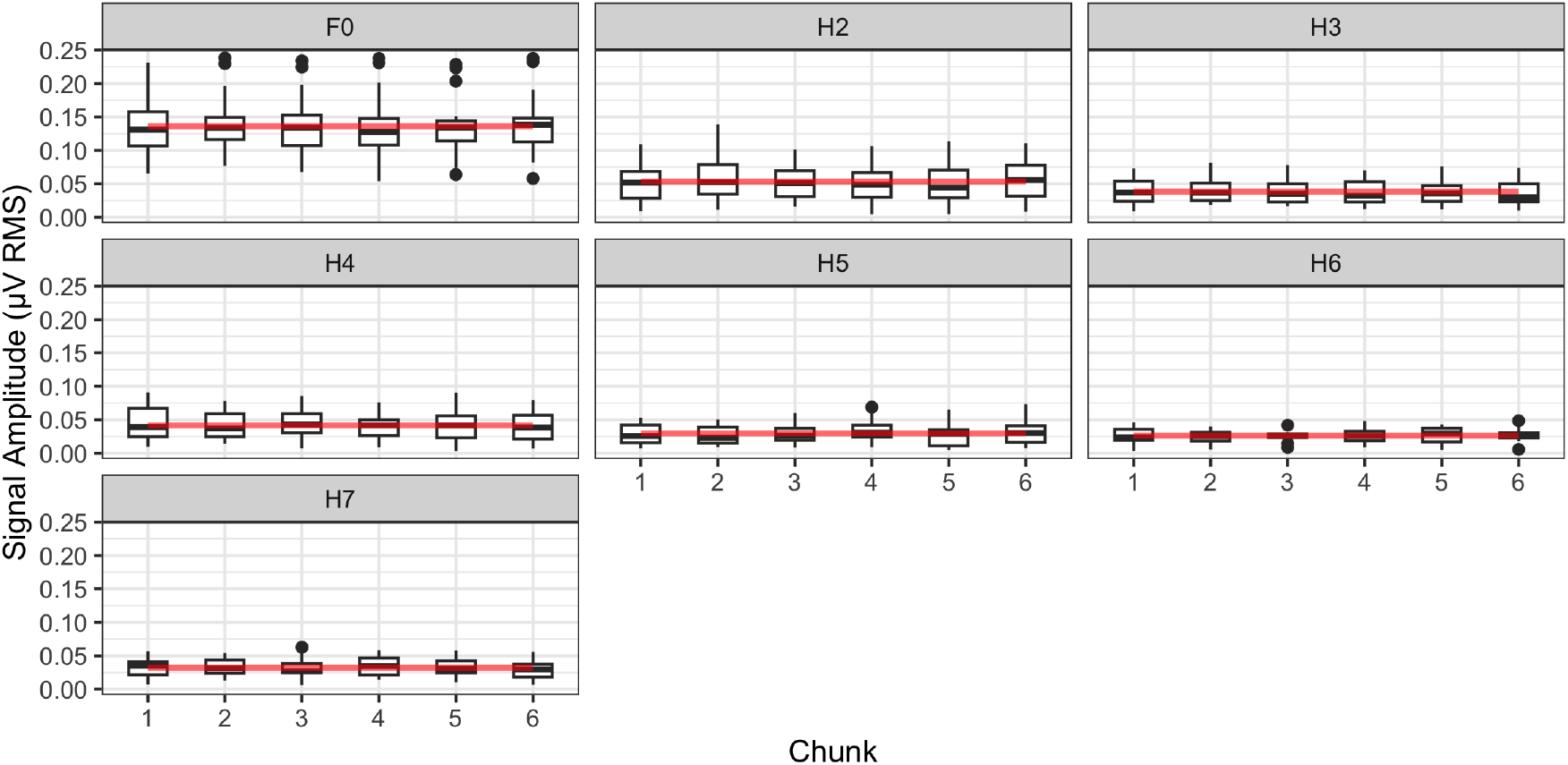
Signal levels of the Rapid FFR over an elapsed time of 9 minutes in each polarity. Here, we isolate 1/6th of the data in sequential chunks. Red lines show the values predicted by the linear mixed effects models.

For the EFR harmonics, SNRs increased significantly across chunks, reaching a peak around chunk 4. Beyond this point, the growth rate declined (e.g., from 1.08 at chunk 1 to -0.51 at chunk 6), consistent with the significant quadratic relationship in the model. However, this trend is not driven by the signal amplitudes contributing to the SNR, as the model for F0-H3 with signal amplitudes did not retain any *Chunk* terms due to their lack of contribution to the statistical model. Similarly, no effect of *Chunk* was observed in the higher four TFS harmonics, for both SNR and signal amplitudes.

## 6 Discussion

In this paper, we investigated thoroughly a rarely used approach to efficiently collect the Frequency-Following Response (FFR), which we term the Rapid FFR. Through a series of analyses, we demonstrate that the Rapid FFR outperforms the Conventional technique, while still capturing the same neural response to a speech-like steady-state sound, a sawtooth wave.

Firstly, we found that the Rapid FFR outperforms the Conventional FFR when both techniques are constrained to the same elapsed time, specifically a two-minute recording window. Our statistical model revealed a significantly higher SNR for the Rapid FFR, with an improvement of approximately 7 dB across harmonics compared to the Conventional FFR. One potential explanation for this difference could involve the onset transients within the Conventional FFR, which may disrupt the establishment of a clear sustained response. However, further analyses indicate that the observed improvement is mainly attributable to the additional recording time made available by removing silent intervals: by eliminating the breaks between stimulus repetitions, the Rapid FFR allows for approximately 47% more data to be recorded within the same time frame. Consequently, the increased data volume enables more effective noise reduction during the averaging process, resulting in elevated SNR values.

Given the differences in presentation and processing methods between the Rapid and Conventional FFR, it is possible that the Rapid FFR measures a distinct electrophysiological response, potentially one with a delayed onset of around 1 second during continuous stimulation. To test this, we examined the correlation between the Rapid FFR and the Conventional FFR responses within each participant. The assumption was that if response strengths correlate across participants, the two methods likely capture the same physiological response. To evaluate this, we computed seven intraclass correlation coefficients (ICCs), one for each harmonic. Our analysis revealed that statistically significant correlations were primarily observed in the lower harmonics. This outcome is likely aided by the fact that the FFR signal is strongest in these harmonics (Tichko & Skoe, 2017), resulting in better signal estimates through higher SNRs and, consequently, higher correlation coefficients. In contrast, the higher harmonics exhibited weaker correlations, likely due to the increased prevalence of noise, which introduces greater variability and thereby reduces the correlation coefficients. Still, these findings suggest that the Rapid and Conventional FFR capture the same neural response, with variability in correlations primarily reflecting systematic noise differences across harmonics.

In the third analysis, we examined the test-retest reliability of the Rapid FFR technique and compared it to that of the Conventional method. The test-retest reliability of the Conventional FFR has been documented (G. M. Bidelman, Pousson, Dugas, & Fehrenbach, 2018), and we anticipated high reliability, particularly as all measurements were conducted within the same experimental session. This eliminated potential sources of variability, such as slight differences in electrode placement. As expected, the Conventional FFR demonstrated high test-retest reliability in the lower harmonics, with particularly strong ICC values. However, reliability declined in the higher harmonics, where ICC values dropped noticeably. This pattern was reflected in the TFS measures, with only two of the four ICC values reaching statistical significance. A similar trend was observed for the Rapid FFR. Reliability was excellent in the lower harmonics, with ICC values reaching up to 0.9. However, in the higher harmonics, ICC values declined, and only one was statistically significant. We additionally performed a supplementary analysis to test whether the Rapid FFR statistically yields higher ICC values than the Conventional FFR for each harmonic. However, no significant differences emerged. Hence, the results suggest that the Rapid FFR achieves comparable test-retest reliability to the Conventional FFR. This finding further supports the Rapid FFR’s validity as a measure of the same neurophysiological process.

In the second experiment, we shifted focus from comparing the Rapid and Conventional FFR to examining the issue of neural adaptation within the Rapid FFR. To investigate this, we presented the same sawtooth wave for 9 minutes in each polarity and analysed if SNR values and/or signal amplitudes changed when examining separate chunks of data across the testing session.

More specifically, we divided our 9-minute recordings into 6 isolated chunks. If neural adaptation was present, we would expect to observe a notable decrease in response strength between the beginning and end of the nine minutes. Our statistical model for SNR values revealed an inverted U-shaped function for the first three harmonics, rather than a linear decline. However, the effect was marginal, with SNR values never deviating by more than 1.8 dB across the six chunks. For the higher harmonics, there was no effect of *Chunk*. Moreover, when analysing signal amplitudes alone in the statistical model, neither the lower nor higher harmonics exhibited significant trends related to *Chunks*.

The negligible impact of neural adaptation may contribute to the ongoing discussion of the generators of the FFR (Glaser et al., 1976, G. M. Bidelman, 2015, Gorina-Careta et al., 2021). Adaptation is less pronounced in lower brain regions, such as auditory nerve fibers, which maintain steady firing rates after an initial decrease over a few 10s of milliseconds (Westerman & Smith, 1984). In contrast, cortical regions exhibit strong adaptation effects, as seen in paradigms like the mismatch negativity. Our results align with the hypothesis that the FFR originates primarily in subcortical structures, such as the inferior colliculus, rather than the cortex.

## 7 Conclusion

In conclusion, the Rapid FFR provides an efficient alternative for collecting FFRs without compromising response quality or reliability. Its robustness to neural adaptation and comparable performance to the Conventional FFR underscore its potential for both clinical and research applications. Moreover, the Rapid FFR offers significant advantages for auditory testing in challenging populations, such as infants, where even minimal time savings are invaluable. Future studies could further investigate its utility across diverse populations and experimental contexts, solidifying its role as an additional tool in auditory neuroscience.

## 8 Acknowledge

The authors would like to thank Steve Nevard for technical support. This work was supported by a PhD studentship grant funded jointly by Action on Hearing Loss and Age UK (grant S19), a PhD studentship grant funded by the Medical Research Council (grant MR/R015759/1, on the MRC Industrial Collaborative Student route), a Pauline Ashley New Investigator Grant funded by Action on Hearing Loss (grant PA08), and by funding from the National Institute of Health Research University College London Hospitals Biomedical Research Centre. There are no conflicts of interest, financial, or otherwise.

The data that support the findings of this study are openly available in OSF at www.osf.io, doi number 10.17605/OSF.IO/Q7STZ.

