## Appendix for "Rapid FFR: A rapid method for obtaining Frequency Following Responses"

### Appendices

#### A ABR measurements

Normal click-ABRs were obtained when wave V had latency values within 5.34 – 6.08 ms, which is ± 3 SDs around the mean for a 100 µs click at 70 dB nHL (=107.6 dB peSPL). The repetition time was kept at 11/s (Picton, Stapells, & Campbell, 1981).

### B Additional Figures

For Experiment 1:

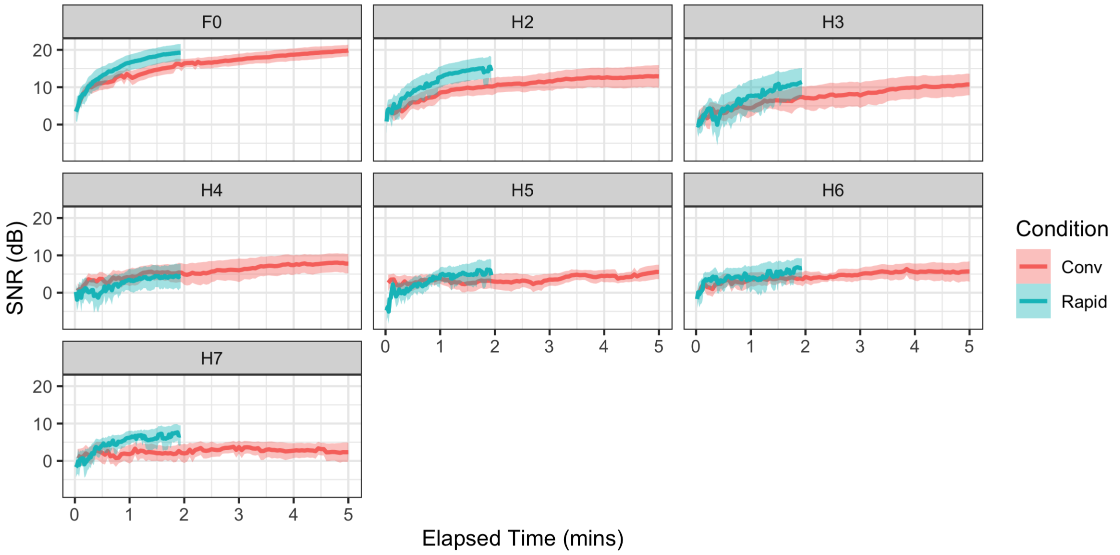


Figure 1: The growth in SNR for the first 7 harmonics of the recorded wave in the rapid and Conventional FFR as a function of elapsed time. Lines represent the mean across participants, and shaded areas show the 95 % Confidence intervals.

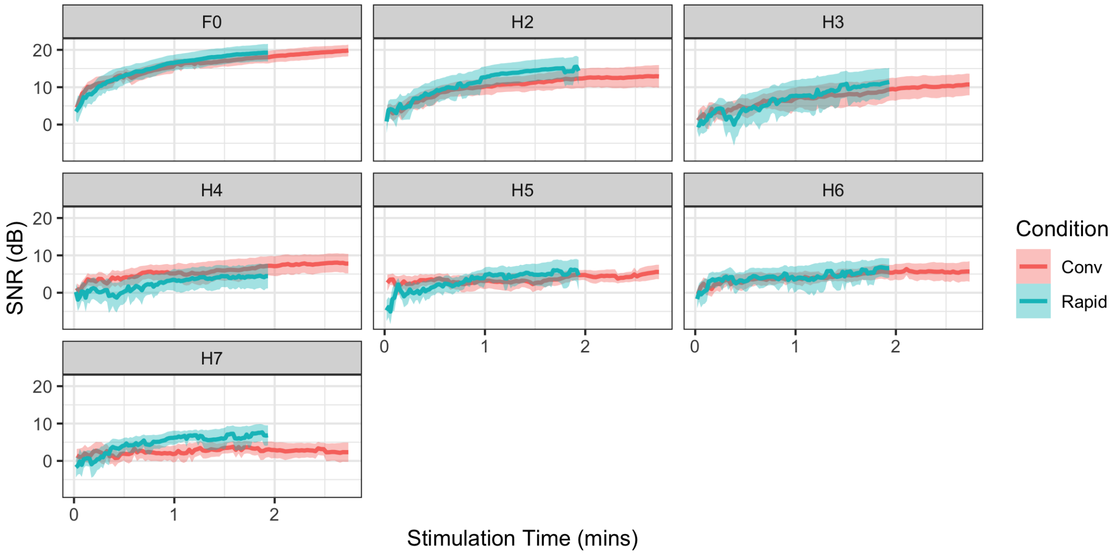


Figure 2: The growth in SNR for the first 7 harmonics of the recorded wave in the Rapid and Conventional FFR as a function of stimulation time. Lines represent the mean across participants, and shaded areas show the 95 % Confidence intervals.

For Experiment 2:

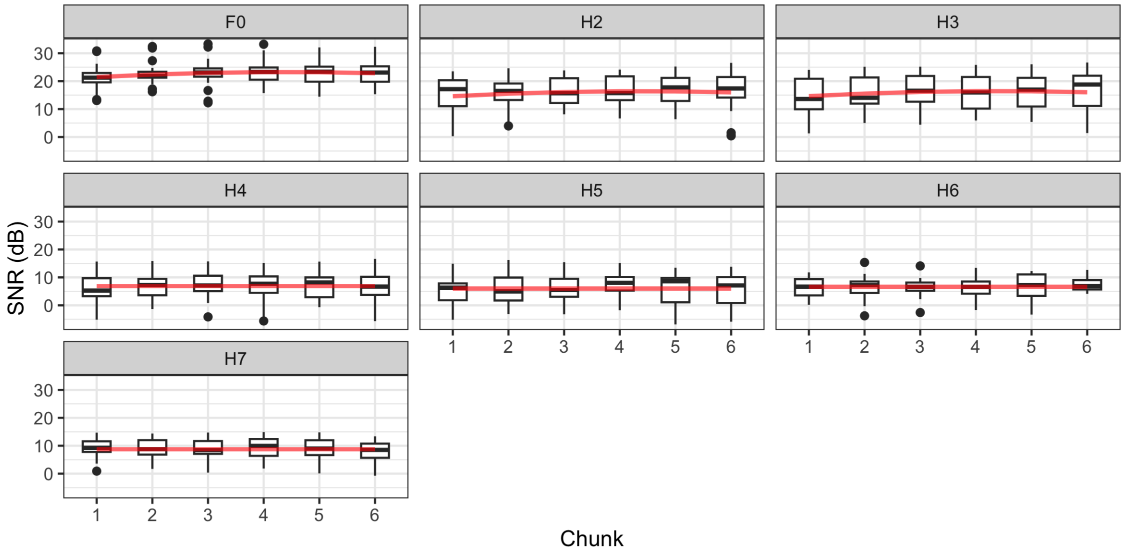


Figure 3: SNR levels of the Rapid FFR over an elapsed time of 9 minutes in each polarity. Here, we independently analyse 1/6th of the data in sequential chunks. Red lines show the values predicted by the linear mixed models.

### C Additional Tables

| factor | Estimate | Std. Error | dF | t-Value | p-Value |
|---|---|---|---|---|---|
| (Intercept) | 20.545 | 1.947 | 222.667 | 10.551 | <.001 * |
| Harmonic | -5.663 | 1.086 | 207.010 | -5.217 | <.001 * |
| I($Harmonic^2$) | .445 | 0.133 | 207.010 | 3.358 | <.001 * |
| ConditionRapid | 6.813 | 2.679 | 207.013 | 2.543 | .001 * |
| Harmonic:ConditionRapid | -2.604 | 1.535 | 207.012 | -1.696 | .091 |
| I($Harmonic^2$):ConditionRapid | .322 | .187 | 207.010 | 1.715 | .088 |

Table 1: Results for the linear mixed model which compares the SNR levels of the Rapid FFR with elapsed time-matched Conventional FFR. Significant p-values here (and in following tables) are marked by a '*'.

| factor | Estimate | Std. Error | dF | t-Value | p-Value |
|---|---|---|---|---|---|
| (Intercept) | 20.545 | 1.786 | 220.742 | 11.503 | <.001 * |
| Harmonic | -5.663 | .996 | 204.989 | -5.686 | <.001 * |
| I($Harmonic^2$) | .445 | .122 | 204.988 | 3.660 | .001 * |
| ConditionRapid | 3.896 | 2.467 | 205.068 | 1.579 | .116 |
| Harmonic:ConditionRapid | -2.276 | 1.416 | 205.100 | -1.608 | .109 |
| I($Harmonic^2$):ConditionRapid | .315 | .173 | 205.137 | 1.821 | .070 |

Table 2: Results for the linear mixed model comparing the SNR levels of the Rapid FFR with data volume-matched Conventional FFR.

| Model | factor | Estimate | Std. Error | dF | t-Value | p-Value |
|---|---|---|---|---|---|---|
| F0-H3 | (Intercept) | 33.959 | 1.878 | 223.804 | 18.087 | <.001 * |
| | Harmonic | -17.181 | 1.623 | 355.034 | -10.589 | <.001 * |
| | I($Harmonic^2$) | 3.443 | .402 | 355.035 | 8.576 | <.001 * |
| | Chunk | 1.389 | .541 | 355.006 | 2.570 | .011 * |
| | I($Chunk^2$) | -0.159 | .076 | 355.002 | -2.099 | .037 * |
| H4-H7 | (Intercept) | 24.923 | 5.666 | 477.744 | 4.399 | <.001 * |
| | Harmonic | -7.453 | 2.122 | 473.263 | -3.513 | <.001 * |
| | I($Harmonic^2$) | .734 | .192 | 473.267 | 3.819 | <.001 * |

Table 3: Results for the linear mixed models which investigates the effect of *Chunk* on the SNR levels of the Rapid FFR.

| Model | factor | Estimate | Std. Error | dF | t-Value | p-Value |
|---|---|---|---|---|---|---|
| F0-H3 | (Intercept) | .287 | .011 | 362.811 | 25.12 | <.001 * |
| | Harmonic | -.184 | .012 | 353.878 | -14.76 | <.001 * |
| | I($Harmonic^2$) | .034 | .003 | 353.894 | 10.94 | <.001 * |
| H4-H7 | (Intercept) | .179 | .019 | 482.2 | 9.278 | <.001 * |
| | Harmonic | -.052 | .007 | 477.4 | -7.212 | <.001 * |
| | I($Harmonic^2$) | .004 | .001 | 477.4 | 6.791 | <.001 * |

Table 4: Results for the linear mixed models which investigates the effect of *Chunk* on the Signal levels of the Rapid FFR.
